# Structural basis for catalysis and substrate specificity of human ACAT1

**DOI:** 10.1101/2020.01.06.896597

**Authors:** Hongwu Qian, Xin Zhao, Renhong Yan, Shuai Gao, Xue Sun, Catherine C. L. Wong, Nieng Yan

## Abstract

Acyl-coenzyme A: cholesterol acyltransferases (ACATs) catalyze acyl transfer from acyl-coenzyme A (CoA) to cholesterol to generate cholesteryl ester, which is the primary form for cellular storage and plasma transport of cholesterol. Because of their close relationship with cholesterol metabolism, ACATs represent potential drug target for the treatment of atherosclerosis and other cholesterol-related disorders. Here we present the cryo-EM structure of human ACAT1 at 3.3 Å resolution for dimer of dimers and 3.0 Å for a dimer. Each protomer consists of nine transmembrane segments that enclose a cytosolic (C) and a transmembrane (T) tunnel. The tunnels, each accommodating an elongated density, converge at the predicted catalytic site. Structure-guided mutational analyses suggest the cytosolic and lateral entry for acyl-CoA and cholesterol, respectively. Our structural, biochemical, and mass spectrometric characterizations reveal the catalytic mechanism and substrate preference for unsaturated acyl chain by ACAT1.

## Introduction

In mammals, cholesteryl ester (CE) is the preferred form of cholesterol for cellular storage and plasma transport (1). Acyl-coenzyme A (CoA): cholesterol acyltransferases (ACATs), also known as sterol O-acyltransferases, catalyze the formation of CE from acyl-CoA and cholesterol (2), a process that is essential for the formation of chylomicron in enterocytes, very low density lipoprotein (VLDL) in hepatocytes, and lipid droplet in adipocytes or macrophages (3). ACATs belong to the membrane-bound O-acyltransferase (MBOAT) family (4). Several key enzymes in lipid metabolism, such as acyl-coenzyme A:diacylglycerol acyltransferase (DGAT) (4) and lysophospholipid acyltransferases (5), and several protein acyltransferases that play a pivotal role in signaling, such as ghrelin O-acyltransferase (GOAT), Wnt acyltransferase (Porcupine), and Hedgehog acyltransferase (HHAT), also belong to this family (4, 6–8).

Two isoforms of ACATs have been identified in mammals, ACAT1 and ACAT2, that share ~ 47% sequence identity (9–12) (Fig. S1). ACAT1 is mostly expressed in liver, adrenal glands, macrophages, and kidneys, while ACAT2 is specifically expressed in intestine and liver (13, 14). ACAT1 is an ER membrane protein consisting of nine predicted transmembrane segments (TMs) (15) and an amino-terminal cytosolic domain (NTD) that is responsible for tetramerization (16). The key active site residue of ACAT1, His460, which is conserved in the MBOAT family, is predicted to locate on TM7 (4, 15, 17). In addition to cholesterol, some steroid molecules, such as sitosterol, cholestanol, allocholesterol, and progesterone, all having the common feature of 3β-hydroxyl group, are also substrates for ACAT1 (18, 19). An extra cholesterol binding site has been suggested for allosteric activation of this enzyme (18–20).

Accumulation of CE in lipid droplets is a main characteristic for foaming of macrophages (21), which may lead to atherosclerotic diseases, the major cause of vascular disease worldwide (22). For the pivotal role of ACATs in converting cholesterol to CE, inhibition of ACATs may be a promising strategy for the treatment of atherosclerosis (23), although the clinical trials remain controversial (24–26). In addition, cholesterol metabolism is closely related to cancer (27) and Alzheimer’s disease (28), the latter as the most common cause of dementia in the western world (29). Using mouse models, inhibitors of ACAT1 have been shown to alleviate amyloid pathology (30), reduce the size of tumors of hepatocellular carcinoma (31), and suppress tumor growth and metastasis of pancreatic cancer (32). Lack of structural information on ACATs has impeded mechanistic elucidation and drug discovery. The only available structure for the MBOAT family is of a bacterial homologue, DltB, which shares low sequence similarity with eukaryotic ACATs (33).

To provide mechanistic insight into ACATs as well as other related MBOAT members, we determined the structure of full-length (FL) human ACAT1 at an overall resolution of 3.3 Å for the dimer of dimers and 3.0 Å for dimer only. We have also established an *in vitro* assay to conveniently measure the activity of ACAT variants. Structure-guided biochemical and mass spectrometric characterizations reveal the substrate entry and product exit paths and elucidate the molecular basis of substrate preference for ACAT1.

## Results

### Structure and enzymatic activity of human ACAT1

The FL human ACAT1, comprising 550 residues, was overexpressed in HEK293F cells with an amino-terminal FLAG tag and a carboxy-terminal 10X His tag and purified to homogeneity through affinity purification followed by size exclusion chromatography (SEC) (Fig. 1A). To examine the enzymatic activity of the purified protein, we established a fluorescence-based assay (Fig. S2A). The purified protein was mixed with cholesterol-containing POPC micelles following a reported protocol (2, 34), and the acyl transfer reaction was initiated by adding oleoyl-CoA into the system. Release of free CoA (CoASH) was detected with a thiol-reactive fluorescence probe, 7-diethylamino-3-(4-maleimidophenyl)-4-methylcoumarin (CPM, Sigma) (35). The catalytic activity measured in the fluorescence-based assay was validated using LC-MS to detect another product, cholesteryl oleate (Fig. 1B).

**Figure 1.**
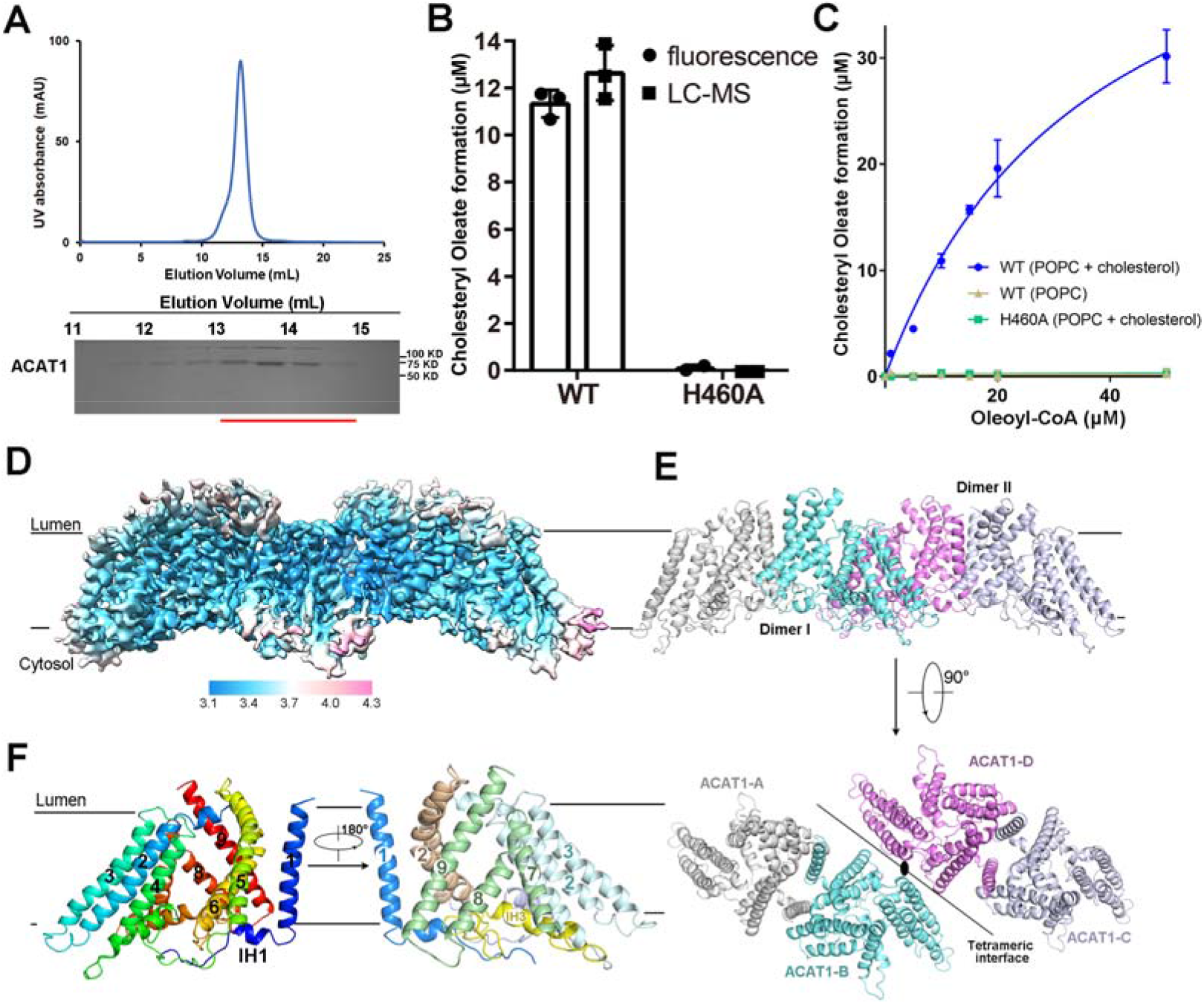
Overall structure of human ACAT1 as a dimer of dimers. **A,** Size-exclusion chromatography purification of ACAT1 using Superose 6 in the presence of GDN. Indicated peak fractions were applied to SDS-PAGE and visualized by Coomassie-blue staining. The highlighted fractions were concentrated for cryo-sample preparation. **B,** Activity assay for purified ACAT1 variants. Two methods were used as cross-validation. Fluorescence: fluorescence-based assay; LC-MS: liquid chromatography–mass spectrometry. Please refer to Methods for details. **C**, Formation of cholesteryl oleate in 10 minutes catalyzed by WT ACAT1 and catalytic mutant (H460A) measured in the fluorescence-based assay. POPC, 1-palmitoyl-2-oleoyl PC. Each data point is the average of three independent experiments and error bars represent S.D. See Methods and Fig. S2A for details of the fluorescence assay. **D,** 3D EM reconstruction of ACAT1. The local resolution map of tetrameric ACAT1, labelled in unit Å, was calculated using RELION 3.0 and prepared in Chimera (45). **E,** ACAT1 exists as dimer of dimers. The two protomers (B and D) that contact each other at the interface between dimers are colored cyan and magenta. Two perpendicular views are shown. The C2 symmetric axis is indicated as an oval dot in the lower panel, and the interface between dimers is indicated by the black line. **F,** Structure of an ACAT1 protomer. The structure is rainbow-colored, blue and red for the amino and carboxyl termini, respectively, on the left, and domain-colored on the right. IH, intracellular helix. All structure figures were prepared with PyMol (46).

ACAT1 purified in 1% (w/v) CHAPS (Anatrace) exhibited better signal to noise ratio than that purified in 0.02% (w/v) glyco-diosgenin (GDN, Anatrace) (Fig. S2B). Therefore, wild-type (WT) ACAT1 and mutants were examined in the presence of CHAPS in all the assays described hereafter. No fluorescence signal was detected in the absence of cholesterol, confirming that acyl-CoA was not hydrolyzed by ACAT1 without cholesterol. Consistent with previous reports, the acyl transfer activity of ACAT1 was abolished when His460 was replaced by Ala (15, 17) (Fig. 1C). A sigmoidal curve (Fig. S2C) was observed with increasing concentrations of cholesterol, consistent with the reported allosteric activation of ACAT1 by cholesterol (34).

It has been shown that ACAT1 prefers oleoyl-CoA to stearoyl-CoA for the transfer activity (34). To elucidate the molecular basis for catalysis and substrate specificity of ACAT1, we set out to determine the structure of ACAT1. Several detergents were tested and the protein purified in 0.02% GDN yielded homogenous particles under cryo conditions.

Details of cryo-EM sample preparation, data collection, and processing are listed in supplemental methods (Fig. S3,4, Table S1). Representative 2D averages indicate an assembly of dimer of dimers (Fig. S3A), consistent with the previous report on the tetrameric organization of ACAT1 (36). After 2D and 3D classification, 348,691 particles were selected to generate a 3D reconstruction at the resolution of 3.3 Å for the dimer of dimers according to the gold-standard Fourier shell correlation (FSC) 0.143 criterion (Fig. 1D, Fig. S3B). To further improve the resolution, we expanded the particles with C2 symmetry and focused on one dimer with an adapted mask for refinement. The resolution was improved to 3.0 Å after further 3D classification (Table S1, Fig. S3C). Atomic model was built based on the dimeric reconstruction (Fig. 1E,F, Figs. S3D, 5).

Although full-length human ACAT1 was used, only 396 residues (118-162, 172-280, 288-440, 443-529) could be resolved in each protomer, among which 391 residues were assigned with side chains (Table S1). In addition to the predicted 9 transmembrane helices (TM1-9) (15), three helices were resolved on the intracellular side, designated IH1, IH2, and IH3, and one on the extracellular side named EH1. While TM1 in each protomer stands out for dimerization, TMs 2-9 are divided into three sub-domains (TM2-4, TM5/6, TM7-9) that are interspersed by two elongated intracellular loops (Loop1 and Loop2), each carrying an intracellular helix, IH2 and IH3, respectively (Fig. 1F, Fig. S3D). The overall transmembrane structure of each ACAT1 protomer appears to be loosely packed with several tunnels and cavities that will be illustrated in detail.

### Dimer of dimers

In the tetrameric ACAT1, the dimer-dimer interface only has limited contact within the membrane, involving TM2/5/6 and IH2 from the two protomers in the center (Fig. 1E, Fig. S6A,B). The loose transmembrane interface supports previous characterization that two putative α-helices in the N-terminal hydrophilic domain (NTD) are required for tetramerization (16).

Corroborating this notion, densities that likely belongs to the NTD can be observed in cluster on the cytosolic side when the map is displayed with low threshold (Fig. S6A). We generated a mutant in which the N-terminal 110 residues were truncated (ΔNTD). The elution volume of this variant appeared about 1.5 mL later than the WT on SEC, indicating disruption of the tetramer (Fig. S6C). To further probe the oligomeric state, cryo-EM images of ACAT1-ΔNTD were acquired and 2D class averages showed reduced size of the particle that is consistent with a dimer (Fig. S6D). These results support that the NTD is responsible for dimerization of dimers, but may not be required for homo-dimerization of ACAT1.

The two protomers in each dimer are similar with root mean square deviation (RMSD) of 1.1 Å over 394 Cα atoms, although no symmetry was applied for data processing (Fig. S6E). The dimer exhibits C2 symmetry around a central axis perpendicular to the membrane plane. TM1, TM5, TM6 and TM9 from the two protomers enclose a deep hydrophobic pocket open to the lumenal side (Fig. S6F,G).

The dimerization of ACAT1 is mainly mediated by extensive van der Waals interactions between TM1 in one protomer and the lumenal segment of TM6 and cytosolic segment of TM9 in the other. In a sense, the two protomers in the dimer swap their respective TM1 segment for tight packing with the rest of the TM domain (Fig. 2). Residues from one protomer, including Val363/Leu364/Phe367/Ile370/Val374 on TM6 and Val501/Trp504/Thr505/Phe508/ Leu509 on TM9, contact Met144/Ile146/ Ala147/Ile150/Leu151/Leu154/Val158 on TM1 from another protomer (labeled as TM1’) (Fig. 2, upper right). On the intracellular side, IH1 also contributes to the dimeric interface. Residues Leu128/Leu132/Val135/His137/Ile138 and Trp408 in each protomers are positioned around the C2 symmetry axis of the dimer to form a hydrophobic interface. In addition, Asp136 and His137 appears to be hydrogen-bonded with Asn409’ and Asp406’, respectively (Fig. 2, lower right).

**Figure 2.**
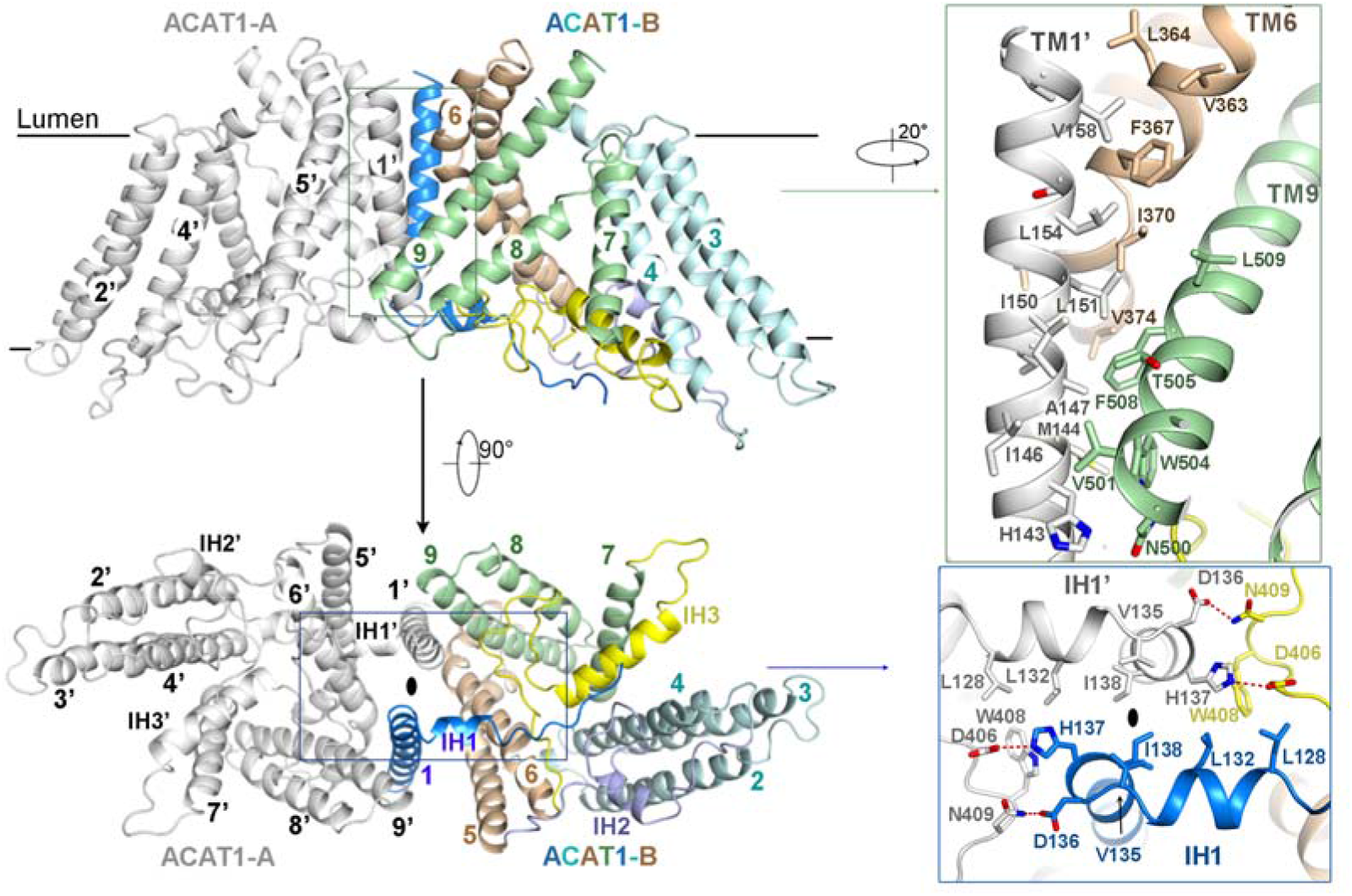
Dimer organization of ACAT1. ACAT1 dimer is mediated by TMs 1/6/9 and IH1 in each protomer. One protomer is colored gray (ACAT1-A) and the other is domain-colored (ACAT1-B). The dimer interface is mediated by the hydrophobic interaction between TM1’ (ACAT1-A) with TM6/9 (ACAT1-B) in TMD (inset, top) and IH1 helices in intracellular side (inset, bottom). The potential hydrogen bonds are represented by red, dashed lines. The C2 symmetric axis is indicated as oval dot.

### A cytosolic tunnel serves as the potential entrance for acyl-CoA

TMs 2-9 of ACAT1 can be superimposed with TMs 3-10 of DltB with RMSD of 5.8 Å over 277 Cα atoms, indicating a conserved catalytic domain in the MBOAT family (33) (Fig. S7A,B). However, there are major conformational differences between ACAT1 and DltB.

In each ACAT protomer, the conserved catalytic residue His460 is located in the middle of the structure, indicating that catalysis may occur within the membrane, with TMs 2-9 enclosing a hydrophilic environment for the acyl transfer reaction (Fig. 3A). TM7 and TM8 constitute a catalytic cavity that opens to the cytosolic side through an elongated tunnel, designated the C tunnel (the cytosolic tunnel). It is noted that the residues that line the C tunnel are conserved in the MBOAT family (Fig. 3A, S1). However, the C tunnel is missing in DltB because of the conformational shifts of TM9, which corresponds to TM8 in ACAT1 (Fig. S7A,C).

**Figure 3.**
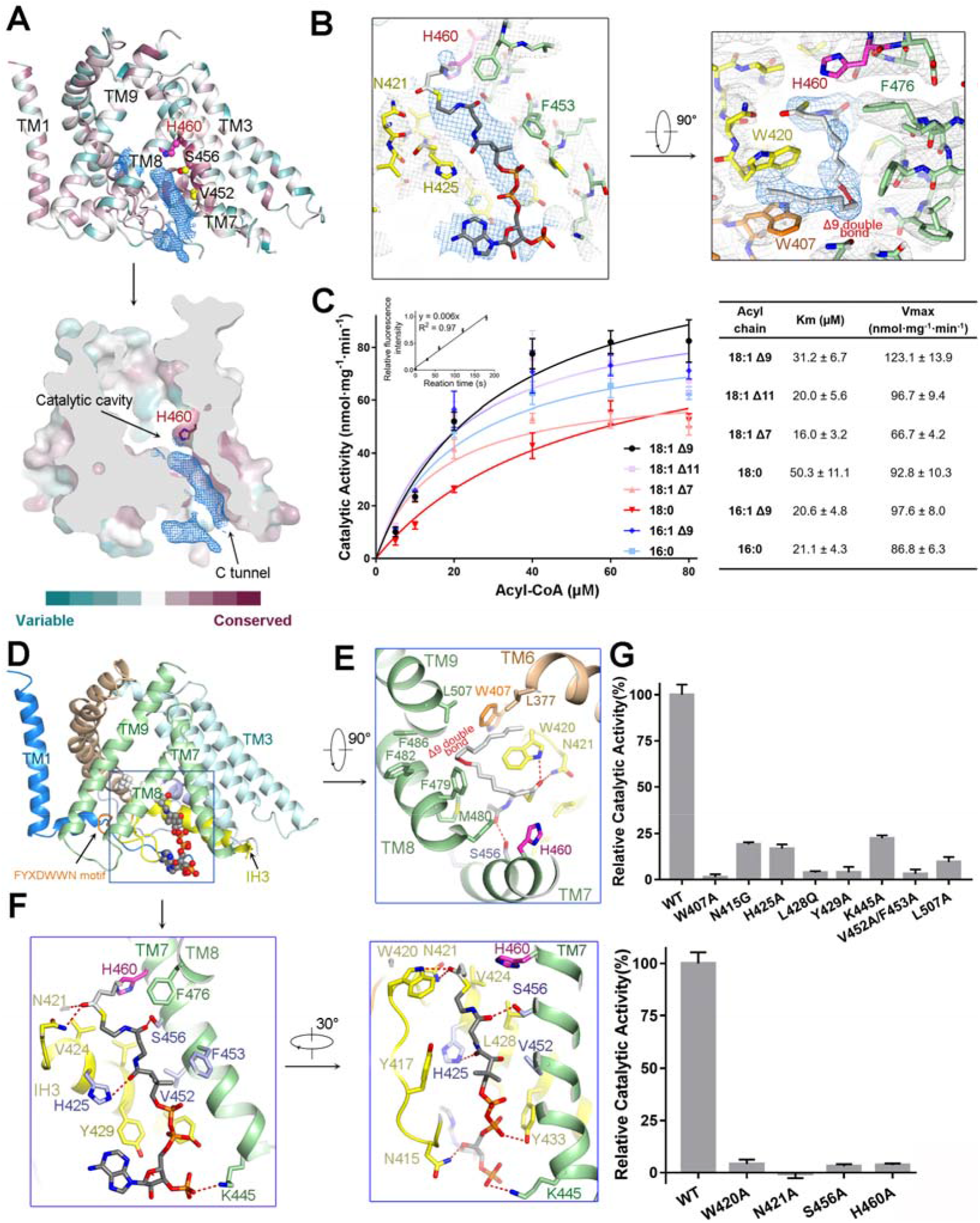
The cytosolic tunnel in the ACAT1 protomer may accommodate oleoyl-CoA. **A,** An elongated density that is reminiscent of oleoyl-CoA is observed in the highly conserved cytosolic tunnel (C tunnel) in each protomer. Structural conservation was analyzed using ConSurf server (47), based on sequence alignment of ACAT1, ACAT2, DGAT1, GOAT, Porcupine, and HHAT from human and mouse using Clustal Omega (48). The density in the cytosolic tunnel, shown as blue mesh, is contoured at 7σ. The mutant containing double point mutations V452Q/S456Q, designated the QQ mutant, was used as negative control in the MS analysis of the bound molecule. His460, Val452, and Ser456 are shown as spheres. See Fig. S8 for the MS results. **B,** Densities of the head group (left) and hydrophobic acyl tail (right) of oleoyl-CoA are contoured at 7σ. Densities shown here are extracted from ACAT1-B, the corresponding densities in ACAT1-A are shown in Fig. S5B. **C,** Substrate preference for ACAT1. *Left*: Enzymatic activity with different substrates. *Right*: K_m_ and V_max_ for substrates acyl-CoA with varied tails. **D**, The oleoyl-CoA binding tunnel is mainly surrounded by TM7/8 and IH3. The FYXDWWN motif is colored orange. **E**, Potential coordination of the oleoyl tail in the catalytic cavity. The contour of the cavity supports preferred substrates to be unsaturated acyl chains. The double bond at Δ9 of the oleoyl tail is colored red. **F,** Potential coordination of the head group of oleoyl-CoA by ACAT1. Oleoyl-CoA is shown as sticks. The catalytic residue His460 is colored magenta. Residues His425, Ser456, Phe453, and Val452, whose biochemical characterizations have been reported, are colored light purple (17, 49). **G,** Functional consolidation of the structurally revealed residues that engage in oleoyl-CoA coordination. The activities of the mutants were normalized relative to that of WT. Please refer to Methods for details.

The bottom of the catalytic cavity in ACAT1 is mainly constituted by the two conserved elongated loops (Loop1 and Loop2), and the top opens to the lumenal side (Figs. S1, 7B,C). Interestingly, an elongated density was observed in the C tunnel of each protomer (Fig. 3A). Because no substrates were added during protein purification or cryo-sample preparation, the density may belong to an endogenous ligand.

Oleoyl-CoA can be best fitted into the density (Fig. 3B). To confirm this analysis, lipids in purified ACAT1 were extracted and applied to liquid chromatography-mass spectrometry (LC-MS) analysis (Fig. S8). Oleoyl-CoA (Sigma), the preferred form of acyl-CoA for ACAT1 (34), was also examined by LC-MS as the positive control. A peak corresponding to oleoyl-CoA appeared in the LC profile (Fig. S6A). The ensuing tandem mass spectrometry (MS/MS) profile of the peak is consistent with the MS/MS fragmentation pattern of oleoyl-CoA, confirming the presence of oleoyl-CoA in the purified enzymes (Fig. S8). Based on structural analysis, a variant designated the QQ mutant (S456Q/V452Q) that is expected to obstruct the cytosolic tunnel was generated to weaken the oleoyl-CoA binding (Fig. 3A). LC-MS of extracts from the QQ mutant showed marked decrease of the oleoyl-CoA peak (Fig. S8A,B), further supporting association of oleoyl-CoA in the purified recombinant ACAT1.

For oleoyl-CoA docking, the head group, including adenosine monophosphate and phosphopantetheine (Ppant), fits reasonably into the density (Fig. 3B, left). At first glance, there was no corresponding density for oleoyl tail contiguous with the head group in the catalytic cavity. But a linear density is found close to the head of oleoyl-CoA. It may belong to the oleoyl tail despite the disconnection from the head density. The oleoyl tail, which is bent at the position of the Δ9 double bond, can be perfectly fit into the density (Fig. 3B, right). Supporting this observation, ACAT1 showed a higher activity for acyl-CoA with the Δ9 double bond than for those with saturated acyl-CoA or unsaturated acyl-CoA with Δ7 or Δ11 double bonds (Fig. 3C).

The oleoyl tail is surrounded by several hydrophobic residues on TM6, TM8, TM9, and Loop2, including Trp420/Trp407 on Loop2 and Leu507 on TM9 (Fig. 3D,E). Ala substitution of Leu507 or Trp407 markedly reduced the catalytic activity (Fig. 3G, top, Fig. S9A). It is noted that Trp407 is positioned in the conserved FYXDWWN motif, which has been suggested to play an important role in acyl-CoA coordination (Fig. 3D,E, Fig. S1A) (37)

The head group of oleoyl-CoA is enclosed by TM7, TM8, IH3 and the loop between TM6 and IH3. The adenine moiety of oleoyl-CoA may be stabilized by the π–π stacking with Tyr429, whose Ala substitution led to a reduction of ~90% enzymatic activity (Fig. 3D,F,G, Fig. S9A). The phosphate appears to be anchored through salt bridge with Lys445, along with hydrogen bonds with Tyr433 and Asn415. Consistent with the structural observation, mutations of Lys445 and Asn415 both led to decrease of enzymatic activity (Fig. 3G, Fig. S9A). The elongated pantetheine arm penetrates the tunnel to place the thioester bond near His460. His425 and Ser456 in the tunnel may form hydrogen bonds with the pantetheine group to properly position the thioester bond at the active site (Fig. 3F). Indeed, H425A reduced the ACAT1 activity by up to 80%, while ACAT1(S456A) only had residual activity, consistent with previous report (17) (Fig. 3G, Fig. S9A).

In addition to the polar interactions, several non-polar residues stabilize the pantetheine group through van der Waals contacts, including Val424/Leu428/Val452/Phe453. Mutational analysis showed that ~ 5% activities remained for ACAT1 variants with the double V452A/F453A or single L428Q point mutations (Fig. 3F,G, Fig. S9A).

### The T tunnel for cholesterol access to the active site in ACAT1

In the structure of ACAT1, a tunnel, enclosed by EH1, TM2/4/5/6, and the Loop 1, traverses the middle of the structure and opens to the center of the membrane. We designate this one as the T tunnel for transverse or transmembrane tunnel (Fig. 4A). Interestingly, the T tunnel and the aforementioned C tunnel converge at the catalytic residue His460. An elongated density is also observed in the T tunnel (Fig. 4A, right, Fig. S10). Although cholesterol cannot be well fit into the density that reaches His460, the T tunnel can perfectly accommodate a cholesterol molecule with the 3β-OH pointing to His460 (Fig. 4B). As the C tunnel appears to be responsible for acyl-CoA entrance and the two tunnels meet at the catalytic site, the T tunnel is thus likely for cholesterol entrance. Supporting the critical role of the T tunnel, Ala substitution of residues near the density, R262A or L306N, resulted in the loss of > 90% activity (Fig. 4C, Fig. S9B).

**Figure 4.**
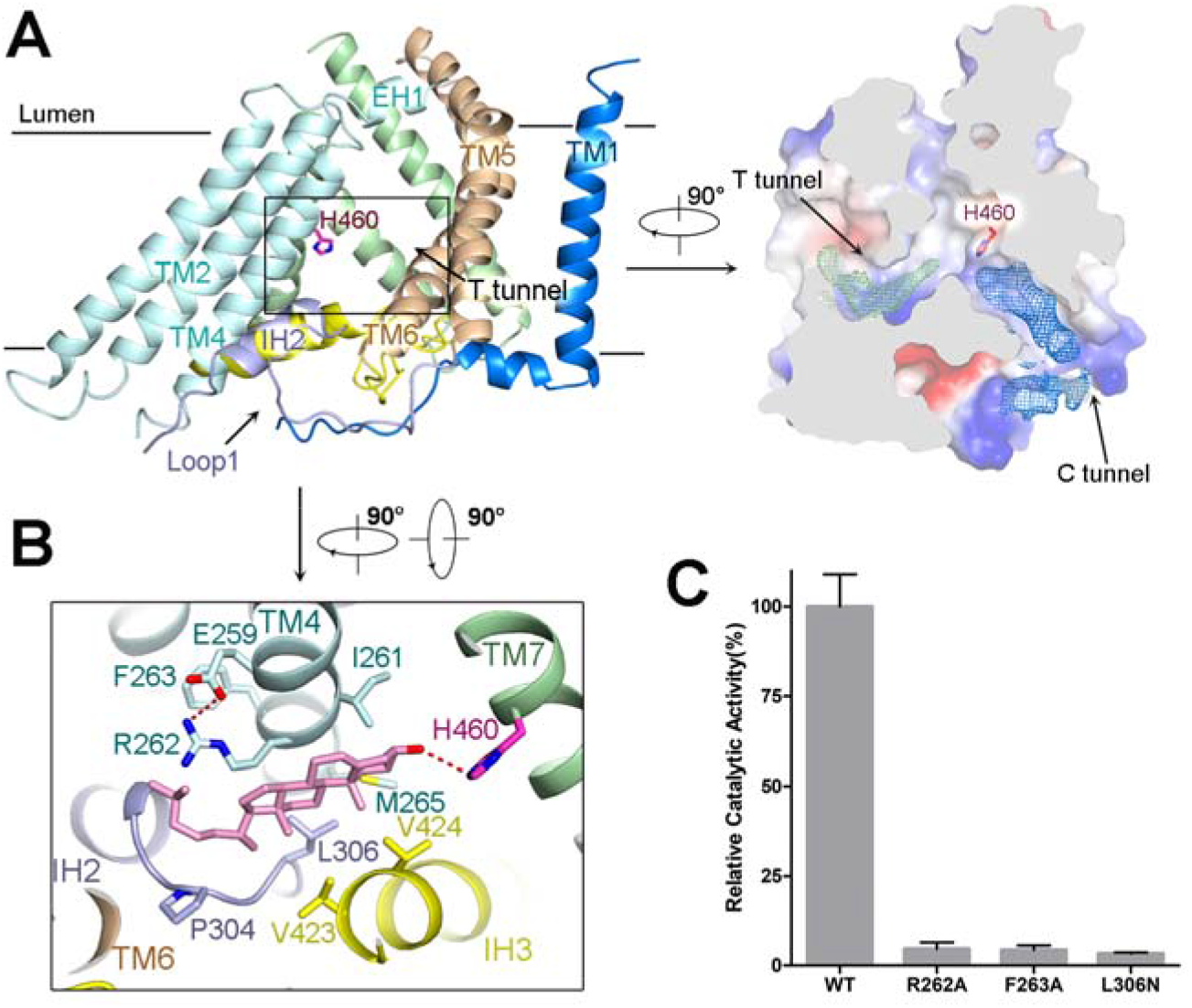
The transverse tunnel may serve as the entrance for cholesterol. **A,** Cholesterol may access to the active site through the transverse tunnel (the T tunnel). *Left*: A side view of one protomer looking through the T-tunnel. The black box indicates the position of the T-tunnel. *Right*: A stretched density is found in the T tunnel. The density, shown as green mesh, is contoured at 7σ. The density, blue mesh, for the potentially bound oleoyl-CoA is also shown at 7σ as a reference. **B,** The T tunnel can accommodate a cholesterol without obvious steric clash. A cholesterol molecule is modelled into the T tunnel with the 3β hydroxyl group within the distance appropriate for hydrogen bonding with His460 (magenta). **C,** Residues constituting the T tunnel are important for ACAT1 activity.

## Discussion

MBOAT members catalyze acyl transfer to the receptors, including lipids and proteins, hence are critical players for lipid metabolism and signal transduction. Based on the structural observation and biochemical analysis, we hereby propose a catalytic model for ACAT1 that will also shed light on the mechanistic understanding of other MBOAT enzymes (Fig. 5).

**Figure 5.**
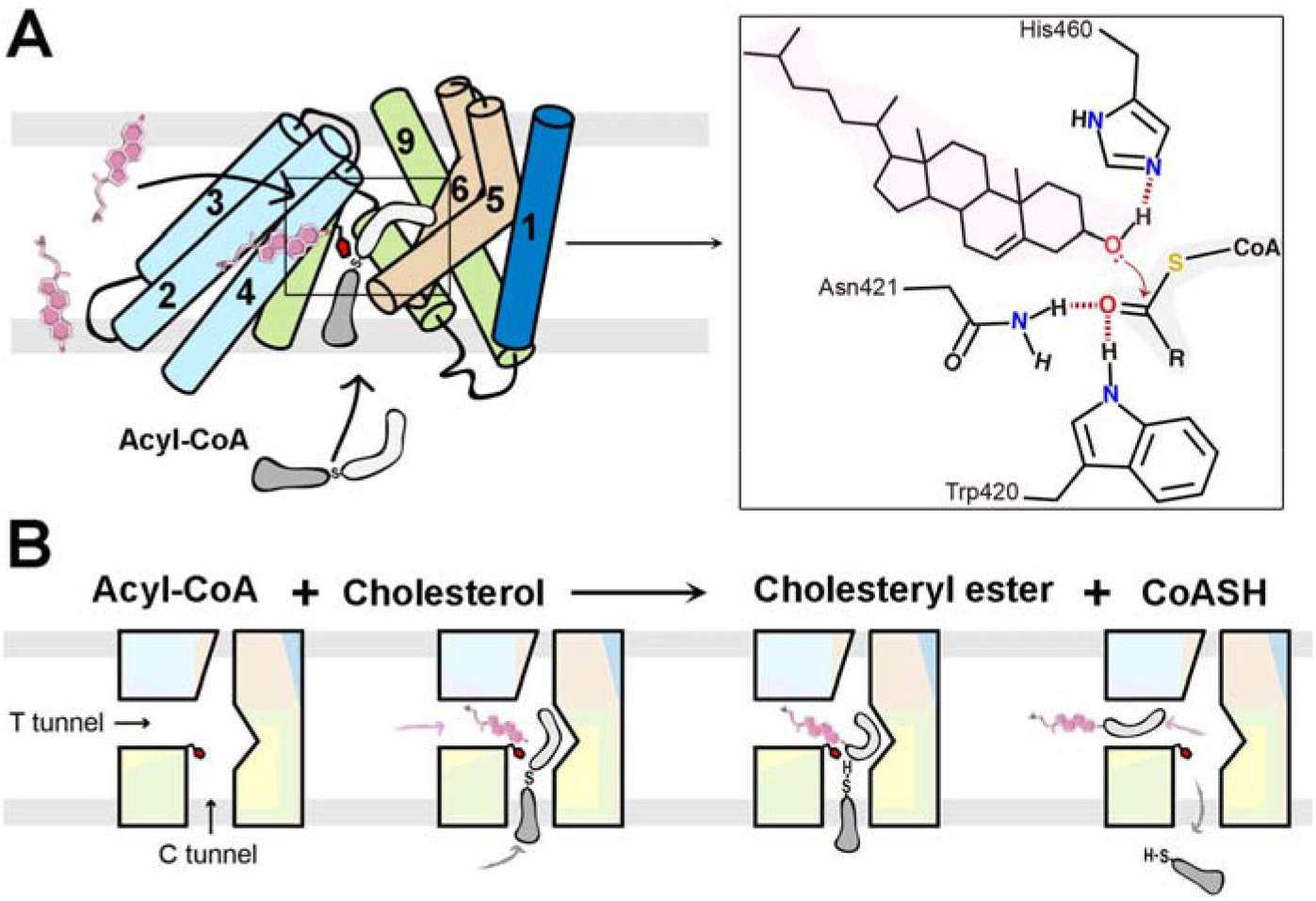
Structure-based working model for ACAT1. **A,** Model for substrate entry and catalysis within ACAT1. The putative mechanism for ACAT1-catalyzed acyl transfer is depicted in the inset: The 3β-OH of cholesterol is activated through deprotonation by His460. The conserved ACAT1 residues Asn421 and Trp420 may stabilize the carbonyl group of acyl-CoA to facilitate the nucleophilic attack. **B,** Working model for ACAT1. The T tunnel and C tunnel may provide the entrance for cholesterol and acyl-CoA, respectively. The reaction is catalyzed in the intersection of the two tunnels, where the catalytic residue His460 is accommodated. The two products, cholesterol ester and CoASH, may exit through the T tunnel and C tunnel, respectively.

The C tunnel and the T tunnel, which converge at the catalytic residue His460, represent the distinct entrance passages for the two substrates, acyl-CoA and cholesterol, respectively. A predicted cholesterol binding motif (306-325) is placed near the entrance to the T tunnel (38, 39) (Fig. S10). Mutations (Y308F and Y312F) in the cholesterol binding motif have been shown to reduce the enzymatic activity (17). Cholesterol may be recruited by the cholesterol binding motif and then enter the active site through the T tunnel.

A His residue is conserved at the active site of several transferases, such as carnitine acetyltransferase (40), cholesterol sulfotransferase (41) and UDP-N-acetylglucosamine acyltransferase (LpxA) (42). The conserved His functions as a general base to deprotonate and activate the nucleophilic substrate (Fig. S11). For acyl transfer catalyzed by ACAT1, the 3β-OH of cholesterol acts as the nucleophilic substrate that is activated through deprotonation, most likely by His460, resulting in the formation of the major product, CE.

In our present structure, the thioester bond of oleoyl-CoA is surrounded by three conserved residues Trp420, Asn421, and His460 (Fig. S1A). His460 and Trp420/Asn421 are on the opposite sides of the thioester of oleoyl-CoA (Fig. 3E,F). His460, highly conserved in MBOAT, has been shown to be pivotal to the catalytic activity (15). Asn421 is also highly conserved and was suggested to act as a second active site (8). Supporting the critical importance of Trp420 and Asn421 for the enzymatic activity, only ~ 5% activity were retained for ACAT1(W420A) and no activity was detected for ACAT1(N421A) (Fig. 3G, Fig. S9A).

In the structure, Trp420 and Asn421 can form hydrogen bonds with the carbonyl oxygen of oleoyl-CoA that may facilitate the nucleophilic attack (Fig. 5A, inset). After exit of CE, which likely diffuses through the T tunnel, His460 is deprotonated with the proton transferred to the sulfur atom to form the other product, CoASH, which can be easily released into the cytosol through the C tunnel (Fig. 5B).

A conserved Ser-His-Asp catalytic triad, including Ser456, His460 and Asp400, was suggested to participate in the catalytic process (17). However, the structure shows that Asp400 is far away from the active site, unlikely to be a component of the catalytic triad. Ser456 is one helical turn below His460 and may help stabilize the latter during catalysis (Fig. S10).

Multiple acyl-CoA molecules can act as the acyl donors recognized by ACAT1, among which unsaturated acyl-CoA molecules are preferred to the corresponding saturated ones. Among the unsaturated acyl-CoA, the contour of the substrate-accommodating tunnel and the enzymatic measurement suggests that Δ9 double bond might be more favored than other positions. Supporting this analysis, mice with mutation in the stearoyl-CoA desaturase 1 (SCD1), also known as Δ9-desaturase that introduces a cis-double bond at the Δ9 position of acyl-CoA mostly stearoyl- and palmitoyl-CoA, were deficient in CE biosynthesis in hepatocytes (43, 44).

Notwithstanding the advances, one major question awaits further investigation, the binding site for cholesterol that serves as the allosteric activator for ACATs (18–20). Nevertheless, the cryo-EM structure of human ACAT1 reported here represents a milestone towards the understanding of the catalytic and substrate selection mechanism of ACAT1 (Fig. 5). The structure and the fluorescence-based assay serve as a foundation for future dissection of the remaining mysteries and for drug discovery.

## Acknowledgments

We thank Paul Shao for technical support during EM image acquisition. We acknowledge the use of Princeton’s Imaging and Analysis Center, which is partially supported by the Princeton Center for Complex Materials, and the National Science Foundation (NSF)-MRSEC program (DMR-1420541). This work was supported in part by the Ara Parseghian Medical Research Foundation (N.Y). H.Q. is supported by the New Jersey Council for Cancer Research. N.Y. is supported by the Shirley M. Tilghman endowed professorship from Princeton University. The atomic coordinates of the tetrameric and dimeric ACAT1 have been deposited in the PDB (http://www.rcsb.org) under the accession codes 6P2P and 6P2J, respectively. The corresponding electron microscopy maps have been deposited in the Electron Microscopy Data Bank (https://www.ebi.ac.uk/pdbe/emdb/) under the accession codes EMD-20239 and EMD-20238, respectively. Correspondence and requests for materials should be addressed to.

## Author Contributions

N.Y., R.Y., and H.Q. conceived the project. H.Q. designed the experiments. H.Q., R.Y., X.Z., X.S and S.G. performed the experiments. All authors contributed to data analysis. N.Y. and H.Q. wrote the manuscript.

