## Supplementary material for "Structural basis for catalysis and substrate specificity of human ACAT1": Materials and Methods, Supplementary Figures S1-S11, Table S1

### Protein expression and purification

The cDNA of human ACAT1 (NCBI reference sequence NM_003101.6) was cloned into the pCAG vector with an amino-terminal FLAG tag and a carboxy-terminal 10X His tag. All mutants were generated with a standard two-step PCR-based strategy. HEK293F suspension cells were cultured in Freestyle 293 medium (Thermo Fisher Scientific) at 37 °C supplied with 5% $CO_2$ and 80% humidity. When cell density reached $2.0\times10^6$ cells per mL, the cells were transiently transfected with the expression plasmids and polyethylenimines (PEIs) (Polysciences). Approximately 1 mg of expression plasmids were pre-mixed with 3 mg PEIs in 50 mL fresh medium and incubated for 15-30 min before transfection. The 50 mL mixture was then added to one-liter cell culture and incubated for 15-30 min. Transfected cells were cultured for 48 h before harvest.

For purification of ACAT1, the HEK293F cells were collected and resuspended in the buffer containing 25 mM Tris pH 8.0, 150 mM NaCl and protease inhibitor cocktails (Amresco). After sonication on ice, the membrane fraction was solubilized at 4 °C for 2 hours with 1% (w/v) GDN (Anatrace). After centrifugation at 20,000 g for 1 h, the supernatant was collected and applied to anti-Flag M2 affinity resin (Sigma). The resin was rinsed with the wash buffer (W1 buffer) containing 25 mM Tris pH 8.0, 150 mM NaCl, and 0.02% GDN. The protein was eluted with W1 buffer plus 0.2 mg/mL FLAG peptide. The eluent was then concentrated and further purified by sizeexclusion chromatography (SEC, Superose® 6 10/300 GL, GE Healthcare) in the buffer containing 25 mM Tris pH 8.0, 150 mM NaCl, and 0.02% GDN. The peak fractions were collected for cryo-sample preparation. The ACAT1 proteins for fluorescence-based assay were purified similarly, except that the buffer for SEC was replaced by 25 mM Hepes pH 7.4, 150 mM NaCl, and 1% CHAPS.

**Fluorescence-based catalytic assay**

To monitor the enzymatic activity of purified ACAT1 and variants, 3 μL 0.8 mg/mL protein was diluted into 37 μL solution containing micelles mixed with taurocholate, cholesterol, and phosphatidyl choline (POPC), as described previously (1), with the final concentration of taurocholate at 4 mM, POPC at 10 mM, and cholesterol at 2 mM in Hepes Buffer (25 mM Hepes 7.4, 150 mM NaCl). To measure the catalytic activity under different cholesterol concentrations, the total amount of POPC and cholesterol was fixed at 12 mM. Another 3 μL of 1M Hepes buffer (pH 7.4) was added to maintain a neutral environment. Acyl-CoA was added to initiate the reaction to the indicated concentration. The mutational analyses were performed in the presence of 40 μM oleoyl-CoA.

Unless specifically mentioned, all the reactions were allowed for 3 minutes at room temperature. The reactions were stopped by adding SDS (Sodium dodecyl sulfate) to a final concentration of 1% (w/v) and then Hepes Buffer was used to dilute the reaction system to 200 μL. For each reaction, 133 μL diluted reaction mixture was transferred to 96 well plate (Greiner Bio-One North America, Inc.), followed by addition of 7 μL 1mM CPM. The mixture was incubated overnight at 4 °C before fluorescence detection using a SpectraMax iD5 Multi-Mode Microplate Reader (excitation, 390 nm; emission, 469 nm). The fluorescence intensity of acyl-CoA free reaction system for WT and each mutant were subtracted to remove the background noise. Nonlinear regression to the Michaelis-Menten equation and allosteric sigmoidal analysis was performed using GraphPad Prism 5.

**LC-MS analysis of cholesterol esterification activity**

The reaction system was the same as described above. Reactions were terminated by adding 200 μL chloroform: methanol (2:1), and in the meantime, 100 ng 18:1 cholesteryl-d7 ester (Avanti) was introduced as internal standard. Samples were vortexed and centrifuged at 5000 rpm for 5 min at room temperature twice. The chloroform layer was collected, dried and resuspended in 40 μL chloroform: methanol (2:1). Serially diluted cholesteryl oleate was used as concentration standard. The cholesteryl oleate was monitored in positive mode by LC-MS using LTQ Orbitrap XL (Thermo Fisher Scientific).

**Cryo-EM sample preparation and data collection**

The cryo grids were prepared using Thermo Fisher Vitrobot Mark IV. Quantifoil R1.2/1.3 Cu grids were glow-discharged with air for 40 s at medium level in Plasma

Cleaner (HARRICK PLASMA, PDC-32G-2). Aliquots of 3.5 µl purified ACAT1, concentrated to approximately 15 mg/mL, were applied to glow-discharged grids. After being blotted with filter paper for 3.5 s, the grids were plunged into liquid ethane cooled with liquid nitrogen. A total of 3,921 micrograph stacks were automatically collected with SerialEM (2) on Titan Krios at 300 kV equipped with K2 Summit direct electron detector (Gatan), Quantum energy filter (Gatan) and Cs corrector (Thermo Fisher), at a nominal magnification of 105,000 × with defocus values from -2.0 µm to -1.2 µm. Each stack was exposed in super-resolution mode for 5.6 s with an exposing time of 0.175 s per frame, resulting in 32 frames per stack. The total dose rate was about 50 $e^-/Å^2$ for each stack. The stacks were motion corrected with MotionCor2 (3) and binned 2 fold, resulting in a pixel size of 1.114 Å/pixel, meanwhile dose weighting was performed (4). The defocus values were estimated with Gctf (5).

**Cryo-EM data processing**

A total of 3,084,959 particles were automatically picked with RELION (6-8). After 2D classification, 1,500,883 particles were selected and subject to a guided multi-reference 3D classification procedure. The references, one good and three bad, were generated with limited particles in advance. A total of 863,672 particles selected from multi-references 3D classification were subject to a global angular search 3D classification with one class and 40 iterations. The outputs of the $30^{th}$-$40^{th}$ iterations were applied to local angular search 3D classification with four classes separately. A total of 650,158 particles were selected by combining the good classes of the local angular search 3D classification. Then a local search multi-reference classification procedure was performed to further classify good particles. In total of 348,691 particles were selected and yielded a reconstruction with an overall resolution of 3.3 Å after 3D auto-refinement with an adapted mask and C2 symmetry. The 650,158 particles were also applied for symmetry expansion using relion_particle_symmetry_expand in RELION (9). After further 3D classification, a total of 760,682 particles were selected to yield a dimeric reconstruction at 3.0 Å after focused refinement with an adapted mask.

All 2D classification, 3D classification, and 3D auto-refinement were performed with RELION 3.0. Resolutions were estimated with the gold-standard Fourier shell correlation 0.143 criterion (10) with high-resolution noise substitution (11).

**Model building and refinement**

The 3.0 Å map for the dimeric form was used for model building of ACAT1. Previously reported crystal structure of DltB (PDB code 6BUI) was used as the initial model to be docked into the map in Chimera (12). Manual adjustment was then made in Coot (13) to generate the final structure. Two dimeric structures were fitted into the tetrameric maps to generate the tetrameric structure with a C2 symmetry. All structure refinement was carried out by PHENIX (14) in real space with secondary structure and geometry restraints.

**LC-MS analysis of lipid extractants from the enzymes**

To prepare samples for LC-MS analysis, 100 μL purified ACAT1 (2.5 mg/mL) in the W1 buffer (25 mM Tris pH 8.0, 150 mM NaCl, and 0.02% GDN) were added into 800 μL extraction buffer (Methanol: Trichloromethane: $H_2O$ = 4:1:3). After centrifugation at 13,000 g for 10 min, the supernatant was transferred to mix with 400 μl 100% methanol followed by 10 min centrifugation at 13,000 g again. The supernatant was dried and re-dissolved in 100 μL LC buffer (methanol/water 7:3) for further LC-MS analysis.

UHPLC (Thermo Vanquish LC system) tandem ESI-Q-orbitrap MS (Thermo Q-Exactive Plus) was employed to analyze 2 μL of lipid extractants, which were performed on ACQUITY UPLC® BEH $C_{18}$ column (50 mm × 2.1 mm, 1.7 μm, Waters, America) with a gradient mobile phase of solvent A (10 mM ammonium acetate in ultra-pure water) and solvent B (methanol) at a flow rate of 0.2 mL/min. The UHPLC elution conditions were optimized as follows: 2% B (0-1 min), linear gradient from 2% to 65% B (1-12 min), 65% B (12-15 min), 65% to 100% B (15-20 min), 100% B (20-25 min), 100% to 2% B (25-28 min) and 2% B (28-30 min). The eluted lipids were directly introduced into MS with vaporiser temperature of 350 °C and spray voltage of 3500 V under negative ionization mode by electro-spray ionization ($ESI^-$). The analytes were monitored under full scan-dd$MS^2$ mode with mass ranged from m/z 100 to m/z 1200. The MS/MS analysis was conducted with collision energy 35 V, resolution 70,000 FWHM, AGC target $5e^4$, maximum IT 200 ms, and isolation window m/z 2.0, respectively. Xcalibur 4.1.50 and Compound Discover 3.0 softwares were utilized to acquire and screen MS and MS/MS data.

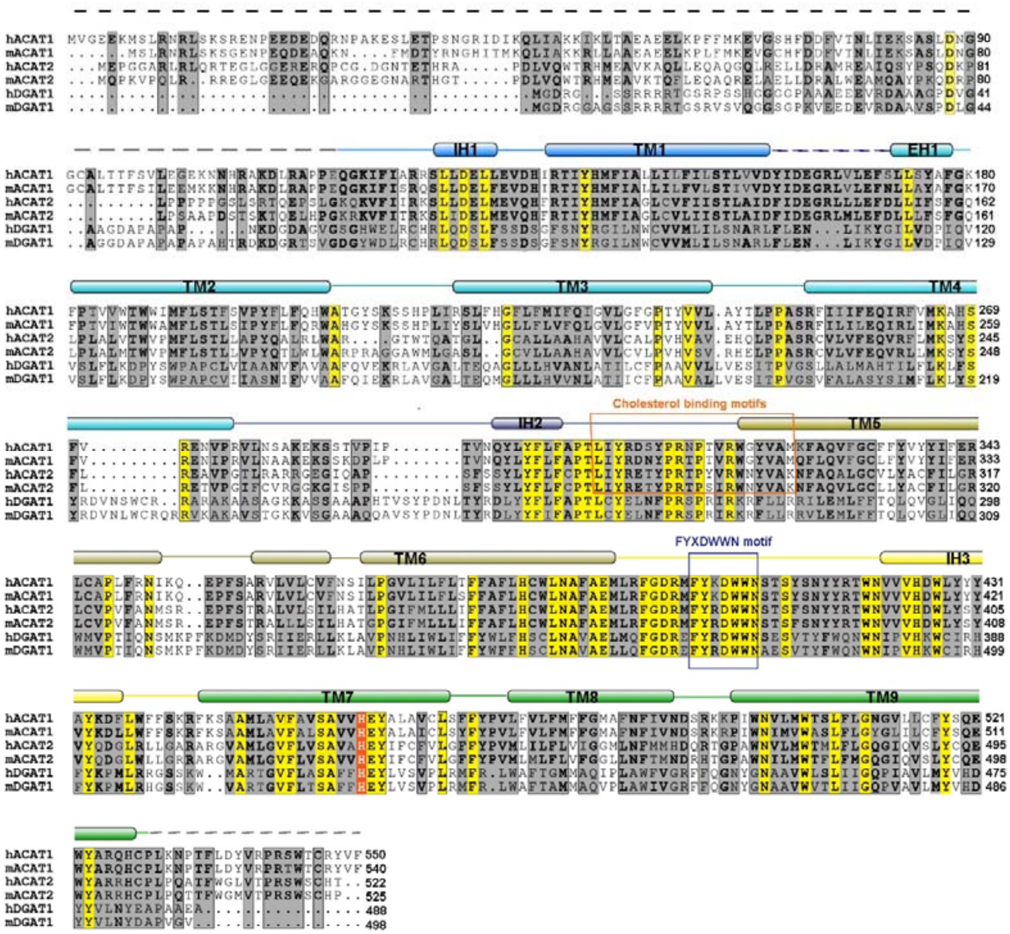


**Figure S1 | Sequence alignment of ACAT1 with ACAT2 and DGAT1 from human and mouse.** Secondary structure elements of human ACAT1 are indicated above the sequences according to the present cryo-EM structure and domain color-coded the same as that for the structure. Invariant and highly conserved residues are shaded yellow and grey, respectively. The conserved residue His460 at the active site is shade red and colored white. The conserved FYXDWWN motif and predicted cholesterol binding motif are indicated with blue and orange boxes, respectively. Sequences from two species, human (h) and mouse (m), were aligned with online MultiAlin server (http://multalin.toulouse.inra.fr).

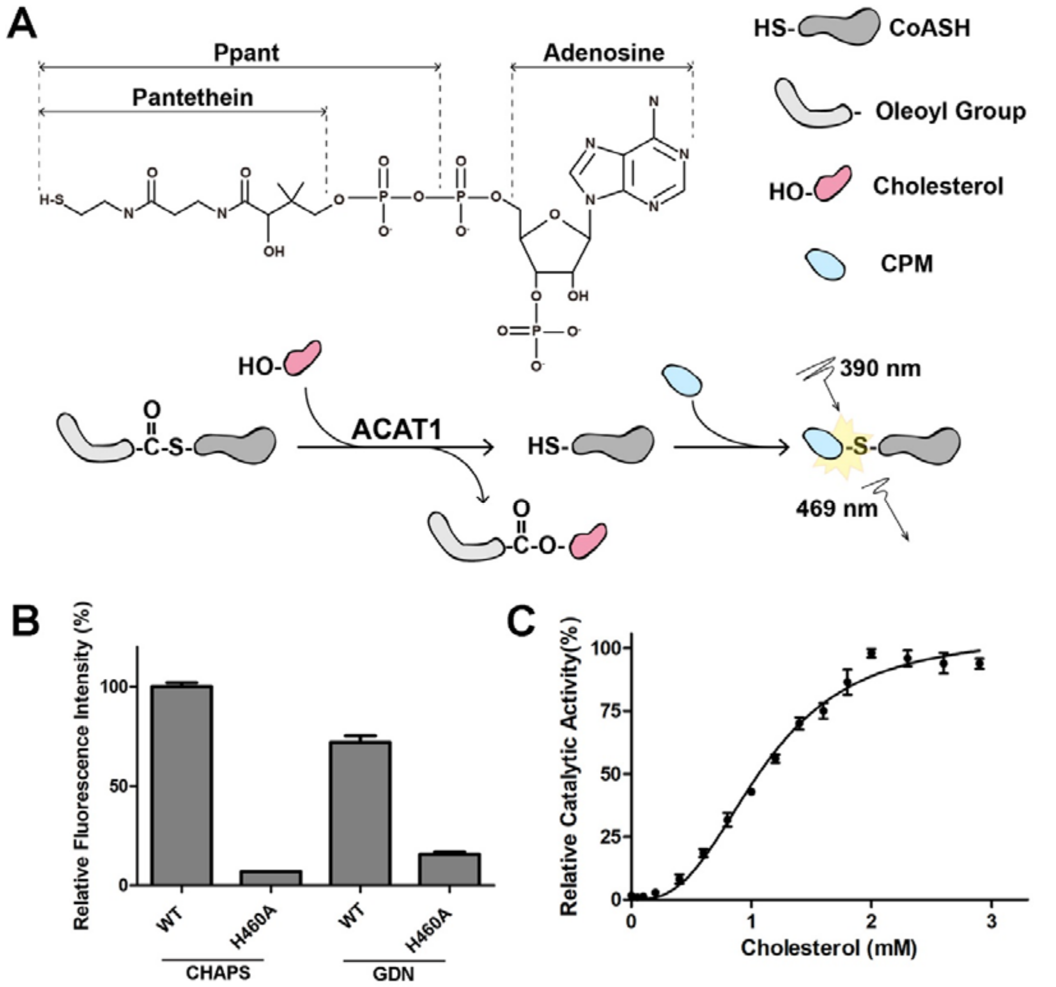


**Figure S2 | Enzymatic activity of recombinant human ACAT1.** **A,** Schematic diagram of the fluorescence-based activity assay for ACAT1. *Top*: the chemical structure of CoASH. *Bottom*: a schematic illustration for the fluorescence-based activity assay. **B,** Effect of detergents on the enzymatic activities of ACAT1. The proteins used for assay were purified from SEC in the presence of 1% CHAPS or 0.02% GDN. **C**, Allosteric activation of ACAT1 by cholesterol. The sigmoidal plot of the catalytic activity with increasing concentrations of cholesterol is consistent with the proposed allosteric activation of ACAT1 by cholesterol (15).

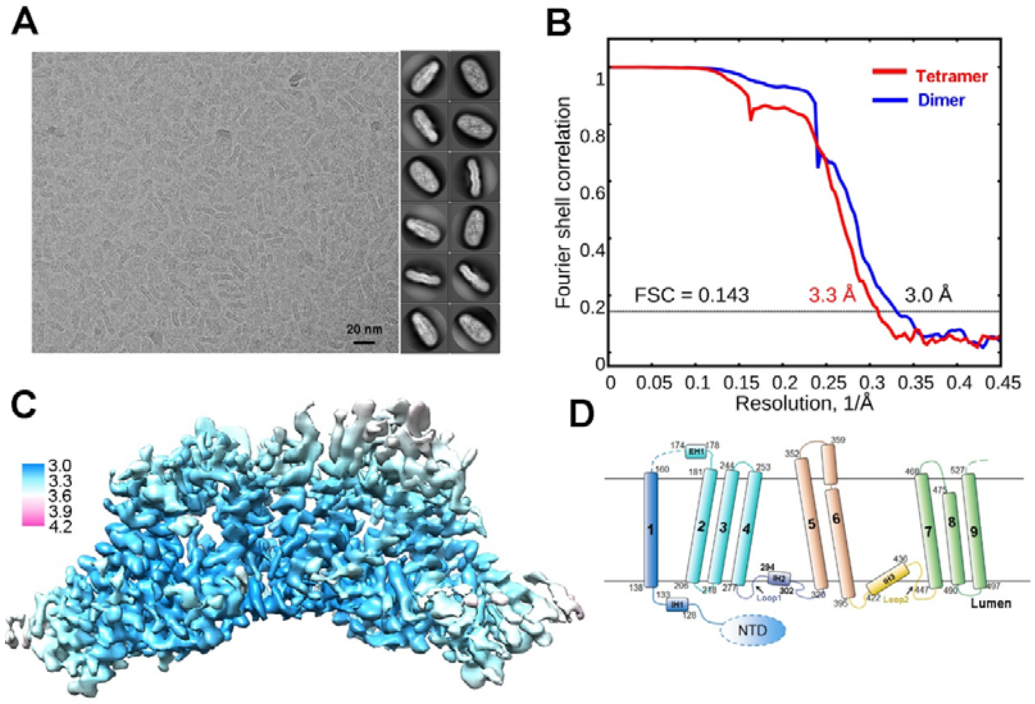


**Figure S3 | Cryo-EM structural analysis of human ACAT1.** **A,** A representative micrograph (left) and 2D class averages (right) of cryo-samples of ACAT1 in GDN micelles. The box size for 2D averages is 310 Å. **B,** FSC curves for the 3D EM reconstructions of tetrameric and dimeric ACAT1. **C,** Local resolution map of dimeric ACAT1 calculated using RELION 3.0. The resolution indicated on the right is shown in unit Å. **D,** Topological structure of ACAT1, color-coded the same as in the right panel of Fig. 1F. IH, intracellular helix; EH, extracellular helix; NTD, N-terminal domain.

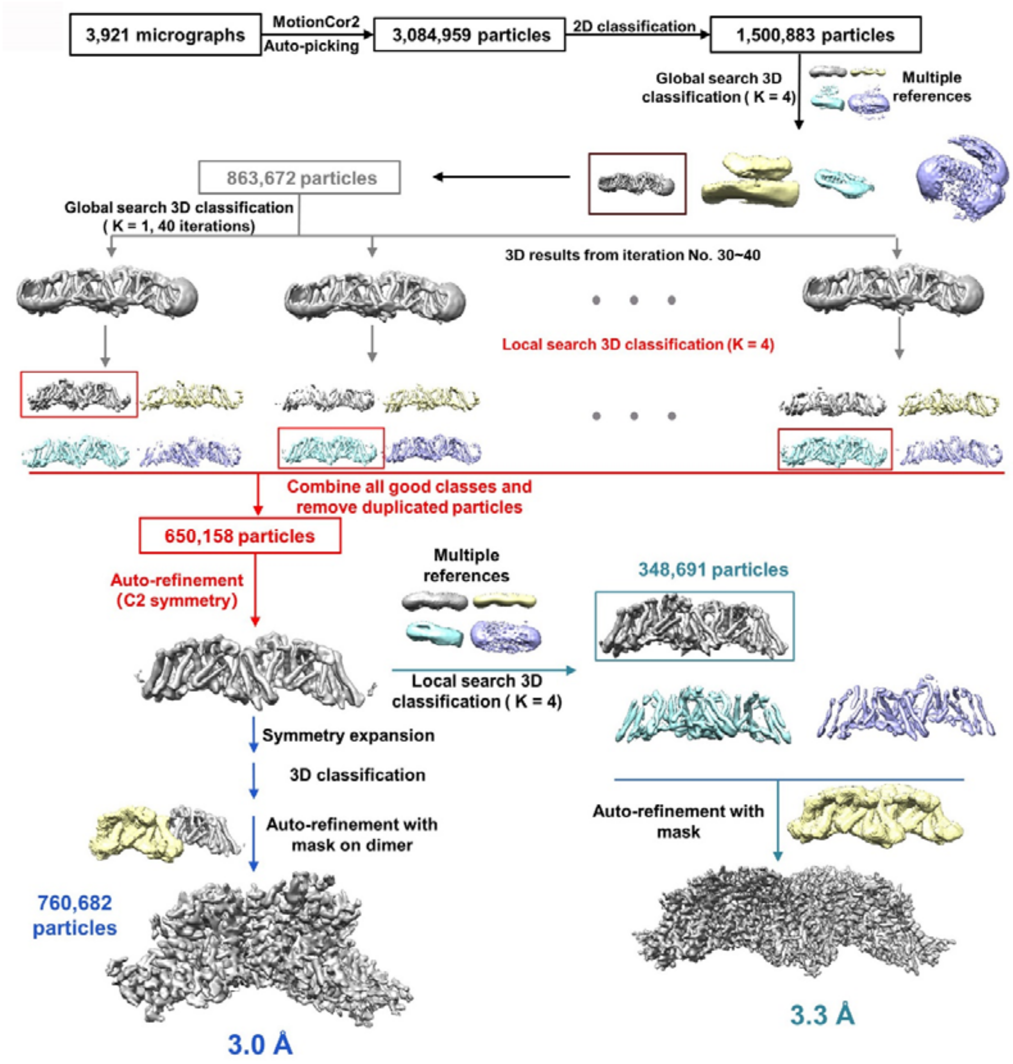


**Figure S4 | Flowchart for data processing.** Details are provided in Materials and Methods.

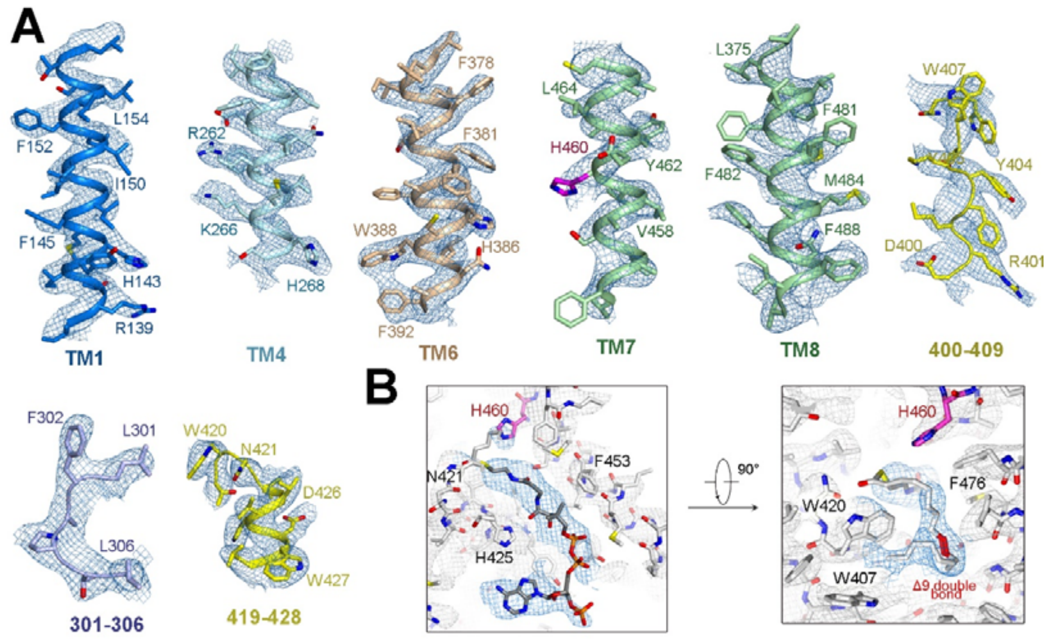


**Figure S5 | EM densities of representative segments.** **A,** The EM maps, contoured at 9-12 σ, were prepared in PyMol. **B**, EM densities for oleoyl-CoA in ACAT1-A protomer, shown as blue mesh, and surrounding residues, shown as grey mesh, are contoured at 7 σ. Two perpendicular views are shown.

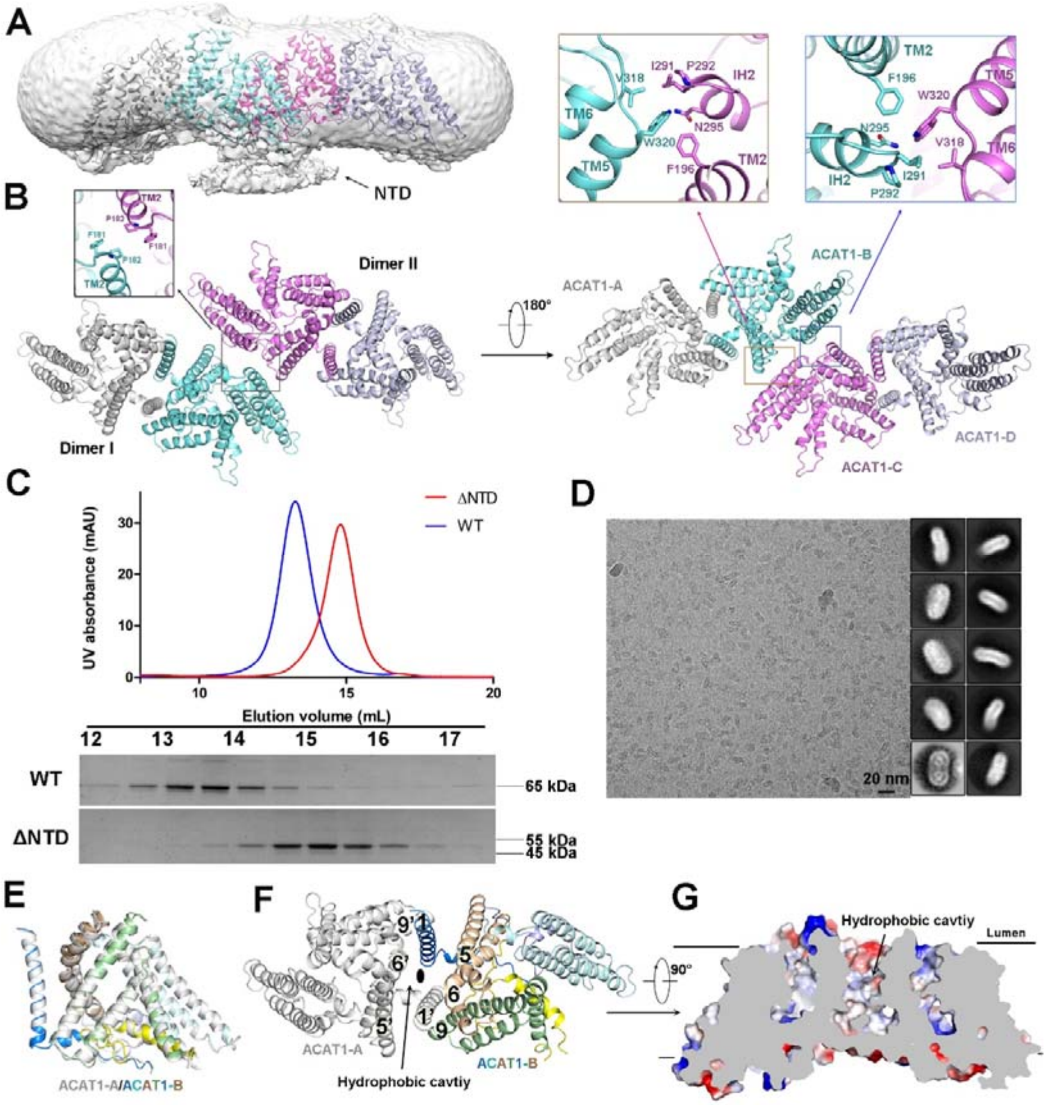


**Figure S6 | NTD is responsible for tetramerization.** **A,** EM map of the tetrameric ACAT1, displayed at low threshold (0.004) in Chimera, reveals extra density on the cytosolic side that may belong to the NTD. **B,** Tetrameric ACAT1 shown in lumenal (left) and cytosolic (right) views, respectively. *Insets*: residues on the tetrameric interface. **C,** NTD is required for tetramerization of ACAT1. Shown here are the SEC profiles and corresponding SDS-PAGE for WT and ΔNTD ACAT1. **D,** A representative micrograph (left) and representative 2D averages (right) of ΔNTD.

Note that the box size for the 2D averages is 220 Å, whereas that for WT ACAT1 is 310 Å. **E,** The two protomers in each dimer are nearly identical. Show here is the superimposition of two protomers, ACAT1-A (grey) and ACAT1-B (domain-colored), in one dimer. **F,** Lumenal view of the dimeric ACAT. An open cavity is formed by TM1/5/6/9 from two protomers around the C2 axis that is indicated by the elliptical dot in the center. **G**, The lumenal cavity in the center of each dimer is highly hydrophobic. Electrostatic surface potential, calculated in PyMol, is shown in side cross section view.

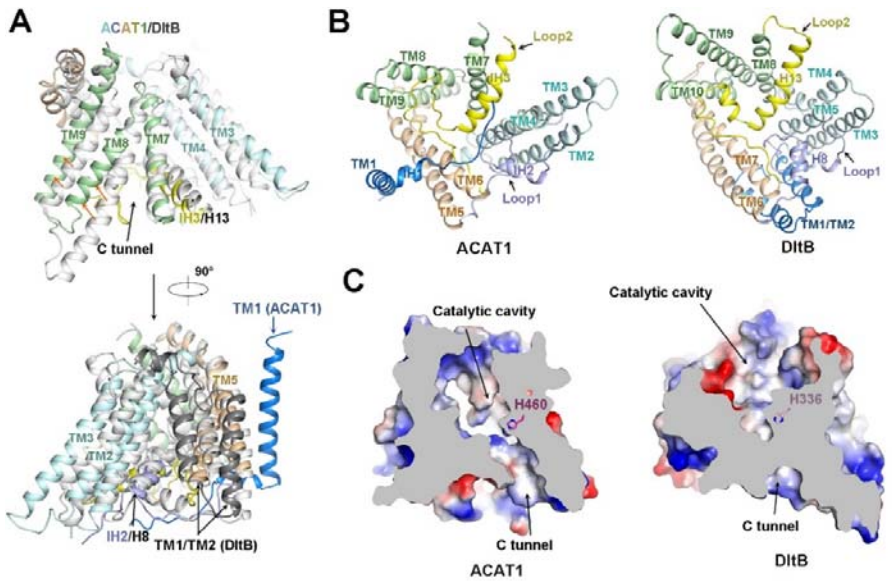


**Figure S7 | Structural comparison of ACAT1 and DltB.** **A,** ACAT1 and DltB have identical structural core. TMs 2-9 of ACAT1 can be superimposed to TMs 3-10 of DltB (PDB code 6BUI) with RMSD of 5.8 Å over 277 Cα atoms. Superimposition of full-length ACAT1 to DltB is shown in two perpendicular side views on the top and bottom. The major conformational shifts of TMs 8/9 of ACAT1 from the corresponding segments in DltB are indicated with orange arrows. TM1 of ACAT1 and the corresponding segments 1/2 (dark) in DltB adopt different structures. **B,** Loop1 and Loop2 constitute are the major cytosolic segments in ACAT1 and DltB. The cytosolic views of ACAT1 and DltB are shown on the left and right, respectively. DltB is domain-colored the same as ACAT1. **C**, The C tunnel can access the catalytic cavity in ACAT1, whereas the corresponding tunnel is closed in DltB. Shown on the left and right are the electrostatic surface potentials of ACAT1 and DltB in the same views. The conserved His is highlighted in magenta in both structures.

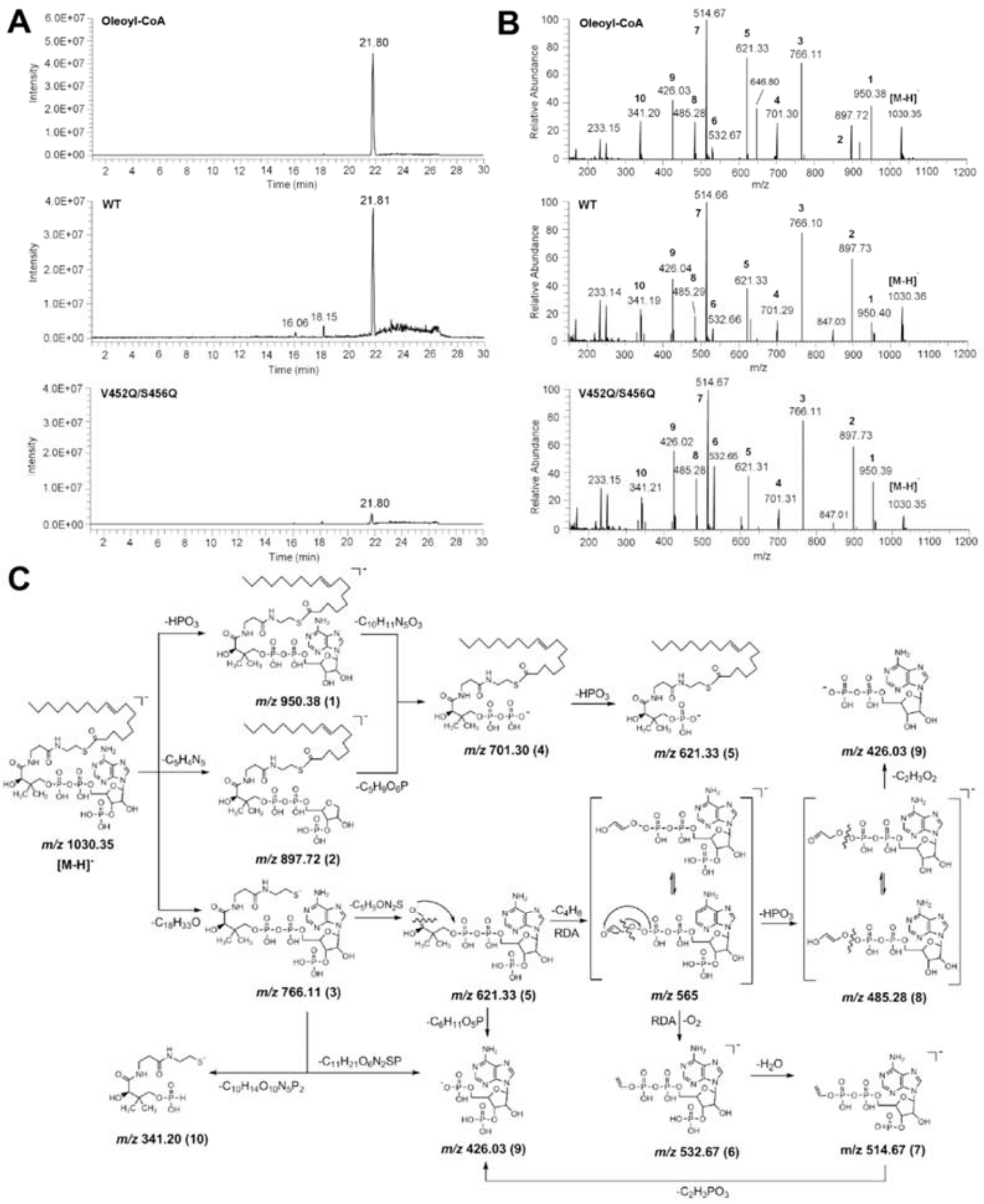


**Figure S8 | LC-MS identification of the ligand to which the linear density in the structure may belong.** **A**, LC profiles of oleoyl-CoA (top), and lipids extracted from WT enzymes (middle) and QQ mutant (bottom). **B**, MS/MS spectrum of purchased (top) and extracted oleoyl-CoA from WT enzymes (middle) and QQ mutant (bottom). Fractions 1-10 represent the same fragments as those in panel C. **C,** Potential MS/MS fragmentation pattern of oleoyl CoA.

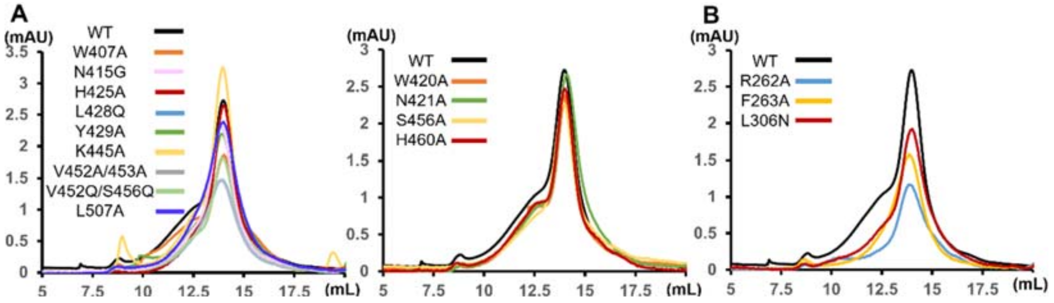


**Figure S9 | SEC profiles of mutants for activity assay.** **A**, SEC profiles of mutants related to oleoyl-CoA coordination. **B**, SEC profiles of mutants for analysis of the T tunnel.

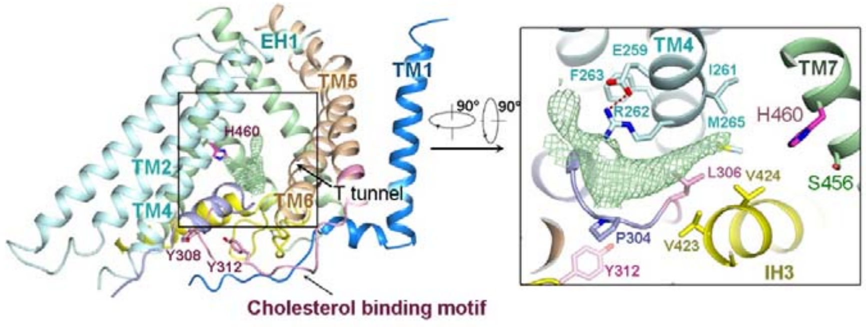


**Figure S10 | The T tunnel for potential cholesterol entrance.** The T tunnel is formed between TM4 and TM6. An elongated density, shown as green mesh and contoured at 7σ, is observed within the tunnel. The predicted cholesterol binding motif is colored pink. *Inset*: lumenal view of the T tunnel in one protomer.

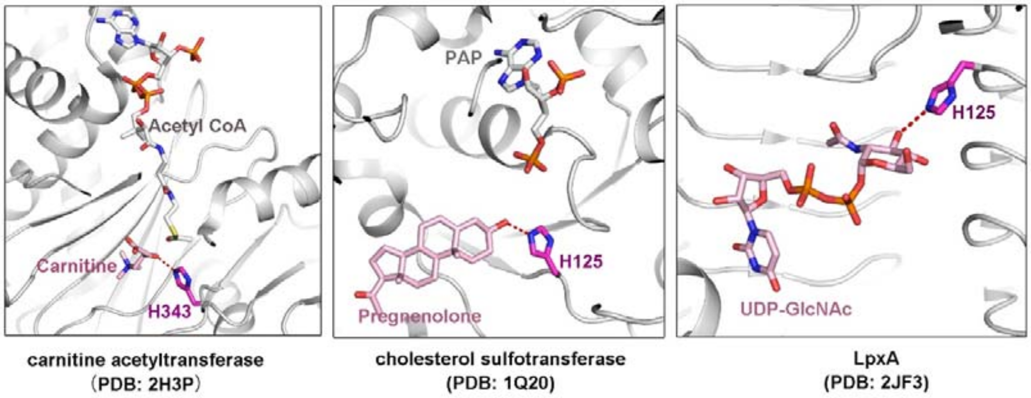


**Figure S11 | Active sites of three representative transferases.** A conserved His residue is found in the active sites in the crystal structures of carnitine acetyltransferase (PDB code: 2H3P), cholesterol sulfotransferase (PDB code: 1Q20), and UDP-N-acetylglucosamine acyltransferase (LpxA) (PDB code: 2JF3). The active His is highlighted in magenta in all three panels. The bound substrates, carnitine, pregnenolone, and UDP-GlcNAc are all colored light pink. The critical His residue may activate the nucleophilic substrate through deprotonation.

**Table S1 | Data collection, 3D reconstruction and model statistics**

| | Tetramer | Dimer |
|---|---|---|
| **Data collection** | | |
| EM | Titan Krios (Thermo Fisher) | |
| Voltage (kV) | 300 | |
| Detector | K2 Summit (Gatan) | |
| Pixel size (Å/pixel) | 1.114 | |
| Electron dose ($e^-$/Å$^2$) | 50 | |
| Number of micrographs | 3,921 | |
| **Reconstruction** | | |
| Software | RELION 3.0 | |
| Number of used Particles | 348,681 | 760,682 |
| Symmetry | C2 | C1 |
| Resolution | 3.3 Å | 3.0 Å |
| Map sharpening B-factor (Å$^2$) | -150 | -104 |
| **Model building** | | |
| Software | Coot | |
| **Refinement** | | |
| Software | Phenix | |
| **Model composition** | | |
| Protein residues | 1584 | 792 |
| Side chain | 1564 | 782 |
| Ligands | 8 | 4 |
| **Validation** | | |
| R.m.s deviations | | |
| Bond length (Å) | 0.01 | 0.004 |
| Bond angle (°) | 1.24 | 0.83 |
| Ramachandran plot statistics (%) | | |
| Preferred | 81.68 | 88.93 |
| Allowed | 17.63 | 10.94 |
| Outlier | 0.69 | 0.13 |
